# High-throughput and genome-scale targeted mutagenesis using CRISPR in a non-model multicellular organism, *Bombyx mori*

**DOI:** 10.1101/2023.08.02.551621

**Authors:** Sanyuan Ma, Tong Zhang, Ruolin Wang, Pan Wang, Yue Liu, Jiasong Chang, Aoming Wang, Xinhui Lan, Le Sun, Hao Sun, Run Shi, Wei Lu, Dan Liu, Na Zhang, Wenbo Hu, Xiaogang Wang, Weiqing Xing, Ling Jia, Qingyou Xia

## Abstract

Large-scale genetic mutant libraries are powerful approaches to interrogating genotype-phenotype correlations and identifying genes responsible for certain environmental stimuli, both of which are the central goal of life science study. We produced the first large-scale CRISPR/Cas9-induced library in a non-model multicellular organism, *Bombyx mori*. We developed a *piggy*Bac-delivered binary genome editing strategy, which can simultaneously meet the requirements of mixed microinjection, efficient multi-purpose genetic operation, and preservation of growth-defect lines. We constructed a single-guide RNA (sgRNA) plasmid library containing 92,917 sgRNAs targeting promoters and exons of 14,645 protein-coding genes, established 1726 transgenic sgRNA lines following microinjection of 66,650 embryos, and generated 300 mutant lines with diverse phenotypic changes. Phenomic characterization of mutant lines identified a large set of genes responsible for visual phenotypic or economically valuable trait changes. Next, we performed pooled context-specific positive screens for tolerance to environmental pollute cadmium exposure, and identified KWMTBOMO12902 as a strong candidate gene for breeding applications in sericulture industry. Collectively, our results provide a novel and versatile approach for functional *B. mori* genomics, and a powerful resource for identifying key candidate genes potential for improving various economic traits. This study also demonstrates the effectiveness, practicality, and convenience of large-scale mutant libraries in other non-model organisms.

## Introduction

In the past centuries, the widely application of model organisms have helped to reveal the basics of molecular biology. However, as the part of the reasons they were chosen for, the current model organisms are too simple and thus cannot answer more complex questions, neither to explain some specific phenomena (Fields and Johnston 2005). With the transition in biology from description to mechanistic understanding, and especially the completion of genome sequencing of massive organisms, more non-model systems are emerging for tackling questions across the whole spectrum of biology, or exploiting the unique biological features of a special organism to address questions of general importance (Goldstein and King 2016). For example, axolotl (Echeverrri et al. 2022), killifish (Beck et al. 2022), volvox (Umen and Herron 2021), ashbya (Wendland and Walther 2005), and diatoms (Jang et al. 2013) have been used to investigate regeneration, aging, evolution, phase transition, and nanobiotechnology, respectively. The silkworm, *Bombyx mori (B. mori)*, is another example, with advantages in demonstrating the biology of Lepidoptera insects, which comprise the vast majority of agricultural and forest pests (Xia et al. 2013; Meng et al. 2017). *B. mori* is also an economically important insect, with sericulture forming a pillar industry in the rural economy of many developing countries.

Currently, a major gap in methodology and resources between the selected few models and other organisms greatly hindering the researchers in choosing and retaining the use of non-model organisms (Russell et al. 2017). In *B. mori*, only a few hundred mutant lines with limited genetic background information have been preserved (Goldsmith et al. 2005, Tong et al. 2022). Functional genomic approaches are lagging, with the first transgenic silkworm obtained in 2000 (Tamura et al. 2000), 17 years after the first transgenic plant (Herrera-Estrella et al. 1983). Widely used RNAi technology has been shown to be extremely inefficient in *B. mori* and many other lepidoptera insects (Terenius et al. 2011; Kolliopoulou and Swevers 2014).

The emergence of CRISPR/Cas9 has expanded genetic toolboxes in a wide range of systems, including *B. mori* (Cong et al. 2013; Jinek et al. 2013; Mali et al. 2013; Ma et al. 2019). Its simplicity and modularity have led to the successful construction of large CRISPR libraries for genome-wide functional genomics and drug target discoveries in mammalian cell systems (Wang et al. 2014; Shalem et al. 2014; Qi et al. 2017; Ford et al. 2019). Large-scale mutant libraries have also been constructed for four model organisms: rice (*Oryza sativa* L.) (Lu et al. 2017; Meng et al. 2017), rapeseed (*Brassica napus*) (He et al. 2023), zebrafish (*Danio rerio*) (Sun et al. 2019) and fruit fly (*Drosophila melanogaster*) (Meltzer et al. 2019; Port et al. 2020; Zirin et al. 2020). CRISPR-based mutants have simple and clear genotype-phenotype correlations, and mutant libraries provide better opportunities to interrogate gene functions than previous libraries, such as those generated through radiation mutagenesis, chemical mutagenesis, random transgenic insertion, or transgenic RNAi (Wenken et al. 2014).

We established a genome-wide CRISPR screen platform in cultured *B. mori* cells and demonstrated its simplicity and efficiency in identifying genes essential for cell viability under normal conditions (Chang et al. 2020a), as well as for screening resistance genes for both biotic and non-biotic stresses (Liu et al. 2021). However, the application of such powerful genome-scale mutagenesis systems at the whole animal level in non-model organisms has been unsuccessful to date. Here, we report the construction of a high-throughput CRISPR/Cas9 mutant library for *B. mori*, a useful resource for silkworm research and breeding. We demonstrate its application for gene function interrogation, as well as potential use for genetic improvement of economic traits. Since *piggy*Bac transposon can drive gene delivery in a wide range of eukaryotes, our strategy and results also lay the foundation for applying genome-scale mutagenesis to other non-model organisms.

## Results

### Design and construction of the CRISPR single-guide RNA (sgRNA) library

Based on our previously reported, highly efficient, CRISPR/Cas9-mediated *B. mori* targeted mutagenesis in the whole animal, and pooled library in cultured cells (Chang et al. 2020a; Ma et al. 2014), we sought to explore the ability of the CRISPR system to generate a library of targeted loss-of-function mutants at whole-animal level. In *B. mori*, genetic transformation is primarily achieved through embryonic microinjection. This process is extremely labor- and equipment-dependent and requires highly skilled technicians; thus, to maximize the efficiency, we designed a high-input/low-output pooled strategy (Fig. 1a). As in the pooled strategy, the target gene of an individual mutant is indicated by the integrated sgRNA sequence itself, we used a binary CRISPR/Cas9 system (Xu et al. 2017) to avoid mis-editing of non-target genes during the early stage of injection. This system expressed Cas9 protein and sgRNA in two separate transgenic lines, and knockout was achieved by crossing them (Fig. 1a).

**Fig. 1.**
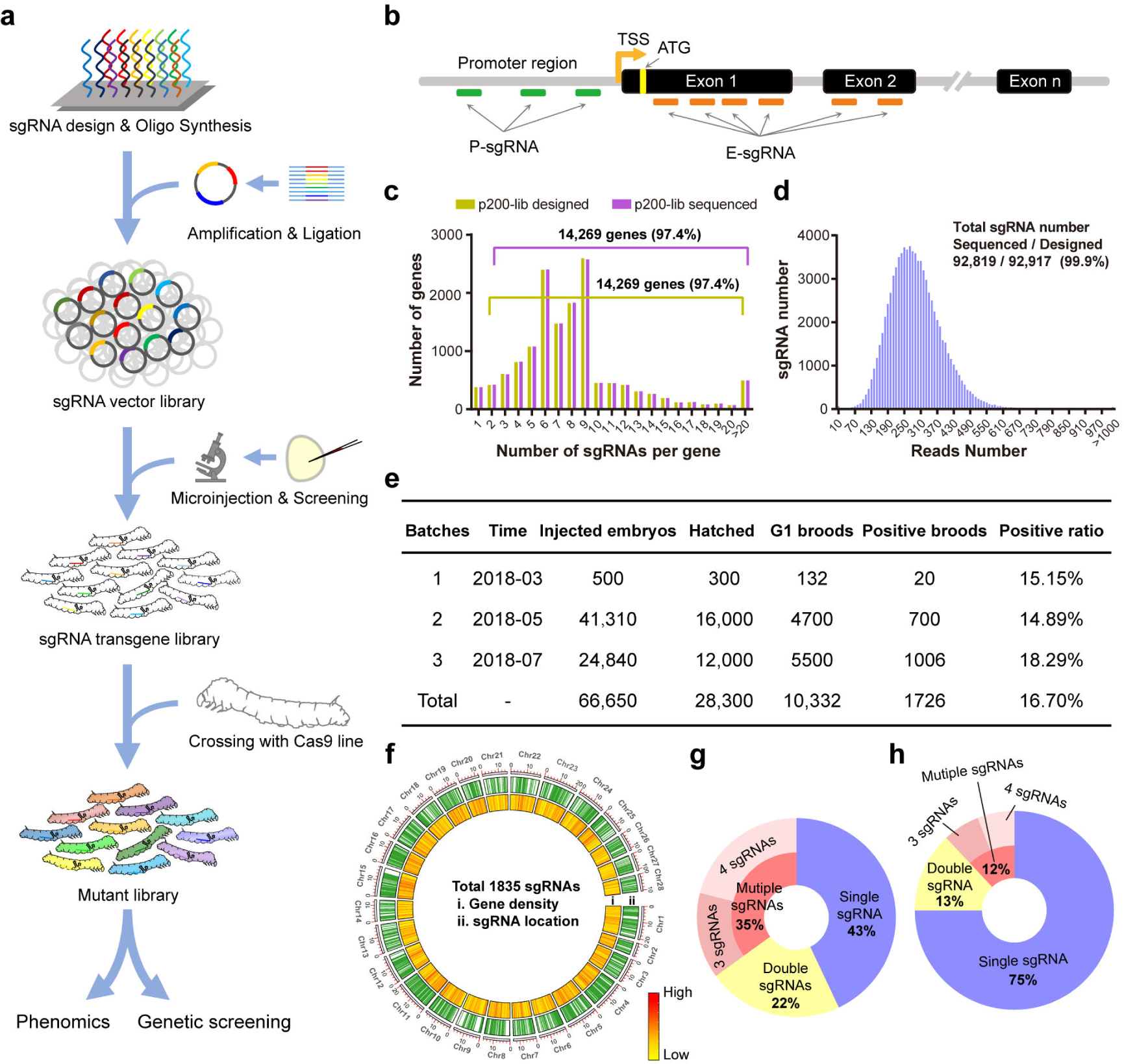
Design and construction of the CRISPR single-guide RNA (sgRNA) library. (a) The strategy of the mutant library construction. (b) The schematic diagram of sgRNA design which target the promoter region and exon around whole genome of B. mori. The sgRNAs target the promoter region were named by P-sgRNA, while sgRNAs target the exon1 or exon2 were named by E-sgRNA. (c) The sgRNA distribution for the p200-lib of designed and deep sequenced. (d) Distribution of the deep sequenced reads and sgRNA numbers. (e) Summary of microinjection information for B. mori. (f) Distribution of sgRNAs detected by deep sequencing over the B. mori. whole genome. (g, h) The number of sgRNAs detected in individual.

To construct the mutant library, we designed a strategy shown in Fig. 1a. The whole genome of *B. mori* was searched for all exons and 500 bp upstream of transcription starting site (Fig. 1b), and 3,489,264 sgRNAs were found (Supplemental Fig. S1). sgRNAs targeting exons were ranked using criteria described previously (Chang et al. 2020a). sgRNAs targeting promoters were ranked using an additional criterion; only sgRNAs targeting the sense strand were selected due to previous findings that the DSBs were repaired in a direction-biased manner (Chang et al. 2020b; Ma et al. 2021). A total of 92,917 sgRNAs were chosen (∼8 E-sgRNAs and ∼2 P-sgRNAs per gene) and synthesized on a microarray chip (Supplemental Table S1). The synthesized sgRNA oligonucleotide pool was elongated by polymerase chain reaction (PCR) and ligated into p200 to form a plasmid library (p200-lib). Deep sequencing of the library showed that 99.9% of the designed sgRNAs were successfully cloned, over 97.4% of the genes had more than one sgRNA (Fig. 1c), and 99.9% of sgRNAs had 36–4109 reads (Fig. 1d). These results indicate sufficient coverage, accuracy, and diversity of p200-lib for subsequent experiments.

### Pooled microinjection and generation of large scale sgRNA transgenic lines

To test the usability and efficiency, a small fraction of p200-lib was co-injected with a *piggy*Bac transposase expression vector (A3-Helper) into 500 G0 embryos. Approximately 300 embryos hatched and were reared to adulthood. Twenty broods positive for 3Xp3-EGFP expression were obtained from 132 G1 broods. The transformation rate was 15.15% (Fig. 1e), which was the average of our previous individual transgenic experiments (Ma et al. 2013). We crossed the 20 transgenic lines with the Cas9 expressing transgenic line (Fig. 1a), and observed clear phenotypic changes in the progenies of the two lines among the 20 F1 lines (Supplemental Fig. S2). We tested the sgRNA of the two lines through simple genomic PCR and found that both phenotypes were theoretically related to their predicted functions (Paine et al. 2019; Schimmel et al. 2021). The target regions of KWMTBOMO05973 and KWMTBOMO01805 were amplified and subjected to deep sequencing. Sequencing results showed that 46% of 12,149alleles and 82% of 22,422 alleles were edited (Supplemental Fig. S2). We concluded that the strategy is suitable for the construction of a genome-wide mutant library, and the established p200-lib plasmid library is suitable for efficient large-scale germline transformation.

Two batches of large-scale micro-injections were performed where 66,150 embryos were injected, 28,000 of which hatched and were reared to adult moths as G1 broods. A total of 1706 broods, containing 1–100 positive transgenic individuals per brood, were obtained through large-scale fluorescent screening of approximately 2,000,000 embryos from 10,200 G1 broods. Average hatchability and transformation ratios were 42.46% and 16.70%, respectively. Given that individuals from the same G1 brood might harbor the same sgRNA, we chose one positive individual from each positive G1 brood and reared them in a pooled manner. Adult moths were crossed back with wild types to generate sgRNA transgenic lines, and an sgRNA library containing 1726 lines was established. To determine the sgRNAs in each line, we performed barcoded deep sequencing and revealed that 1726 lines contain a total of 1835 sgRNAs targeting 1615 genes (Fig. 1f). Over half of the individuals harbored more than one sgRNA, indicating multicopy *piggy*Bac-mediated transformation (Fig. 1g). To generate as many single sgRNA lines as possible, we performed two continuous rounds of backcrossing of these sgRNA lines with wild types and increased the percentage of single sgRNA lines from 43% to 75% (Fig. 1h).

### Genotypic characterization and editing outcome of the mutant library

After generating 1726 sgRNA lines, we investigated the potential of these lines to generate a genome-edited mutant library. As a proof-of-principle demonstration, we randomly selected 300 sgRNA lines and crossed them with Cas9 expressing lines. A total of 150,000 hybrid progenies were reared on fresh food and screened under fluorescent microscopy for mutants. Individuals with both 3Xp3-EGFP and IE2-EGFP positive signals were considered as mutants. To determine editing efficiencies and patterns, we randomly choose 3–6 mutant silkworms from each line and performed deep sequencing of the amplified target regions. Of the 1436 tested silkworms, 89.8% showed detectable editing events (Supplemental Fig. S3), with over 64% showing editing efficiency higher than 50% (Fig. 2a). Such efficiency is sufficient for functional investigation of target genes, and the results encouraged us to improve it. We analyzed the potential parameters that may affect editing efficiency. The GC content was higher in positive than in negative sgRNAs (Fig. 2b). In positive sgRNAs, GC content was 54%, 49%, and 48% in the high-, medium-, and low-efficiency sgRNA groups (Fig. 2c), respectively, with this tendency more significant in the 2–10 nt of the sgRNA (distal to the PAM) region (Fig. 2d), indicating a positive contribution of GC content to editing efficiency. We also found that A/C within the PAM sequences (Fig. 2e) and G at the 3′ end (Supplemental Fig. S4), G at the 5′ end (Supplemental Fig. S4), G/CG/C dinucleotides (Supplemental Fig. S5) within the seed region, and 6mA methylation at the target site (Fig. 2f) are beneficial for efficient editing, whereas A at the 3′ end (Supplemental Fig. S4a) and C at the 5′ end (Supplemental Fig. S4b), and TA/T dinucleotides within the seed region (Supplemental Fig. S5) had a negative effect. We analyzed editing efficiencies of multiple sgRNAs in a single animal and found that the integrated multiple sgRNAs did not affect each other (Supplemental Fig. S6).

**Fig. 2.**
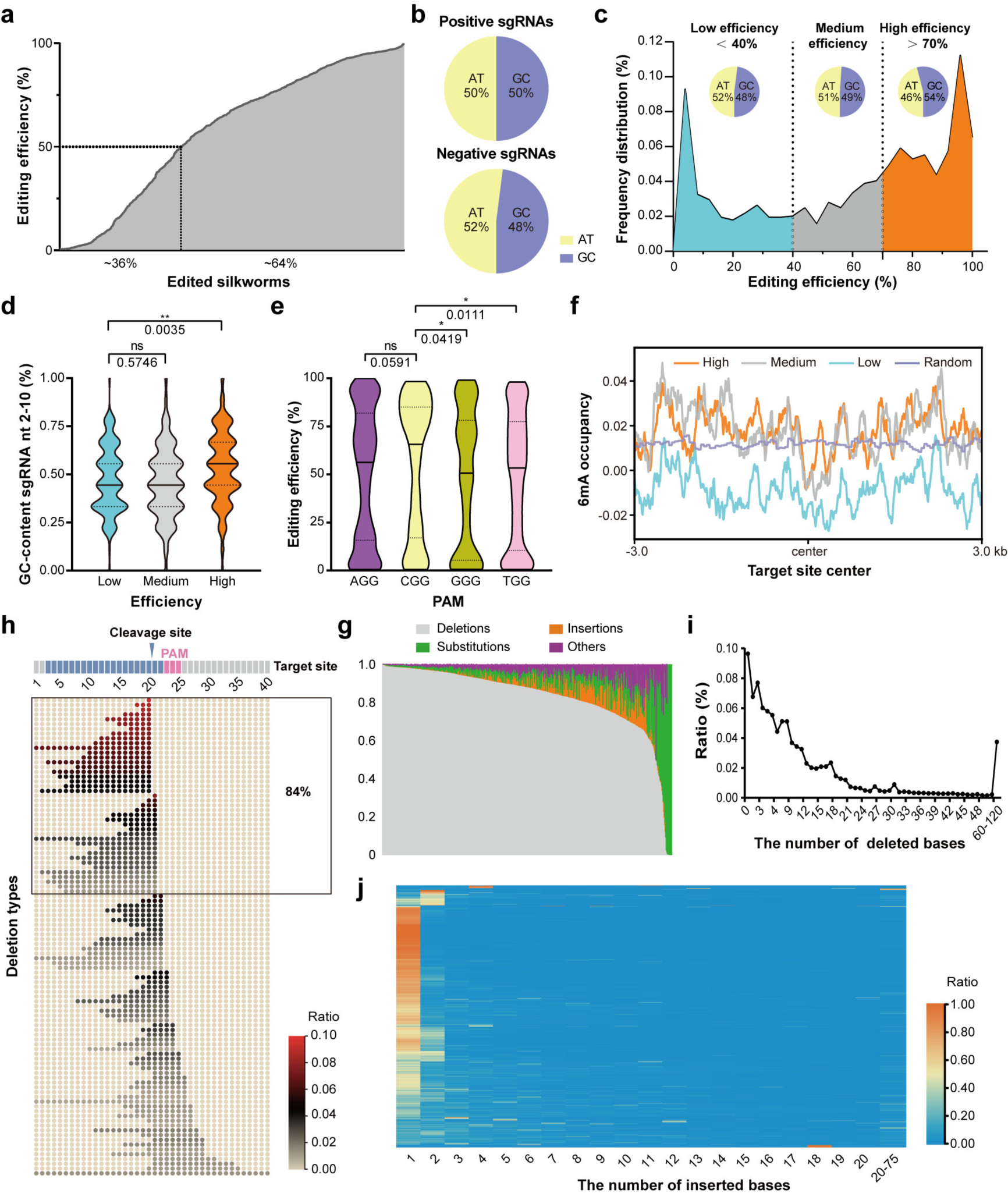
Genotypic characterization and editing outcome of the mutant library. (a) The overall editing efficiency of the edited silkworms. (b) The GC content in the positive and negative single-guide RNAs (sgRNAs). (c) Frequency of distribution of editing efficiency and the GC content in high (editing efficiency > 70%)-medium- and low (editing efficiency < 40%)-efficiency sgRNA groups. (d) The GC content of different editing efficiency. (e) The editing efficiency of different PAM sequences. (f) The DNA 6mA methylation occupancy around the target center with different editing efficiencies. (g) The overall editing patterns of the edited silkworms. (h) The top 100 deletion types and their percentages. (i) The size of the deletions and their occurrences. (j) The size of the insertions and their percentage.

Large-scale deep sequencing of target sites provided the opportunity to investigate editing outcomes in vivo, the patterns of which were previously found to display distinct differences between cultured cells of *B. mori* and that of other species^36,37^. Of the total 61,720, 266 reads, we detected 28,927,046 mutated reads containing 1,387,421 mutation types. Most target sites showed editing patterns mainly composed of deletions (86.50%), with a small fraction of insertions (7.39%) and substitutions (3.28%), similar to that of previous observations (Fig. 2g). A few target sites showed a preponderance of insertion or substitution, while deletion accounted for only a small part (Fig. 2g). More than 84% of the deletions occurred upstream of the cleavage sites (Fig. 2h), which further confirmed the distinct unidirectional feature of DSB repair in *B. mori*. Deletion sizes were generally negatively correlated with their occurrence, with significant occurrence peaks at each 3X base pair (Fig. 2i), indicating a potential beneficial selection mechanism during DSB repair in *B. mori*. Within the insertions, 56.66% and 14.14% were 1 bp and 2 bp insertions, respectively (Fig. 2j). Most insertions were from templated repair, 75% of the 1 bp insertions were duplications of −4 nucleotides of the sgRNA sequence, and 80% of the 2 bp insertions were duplications of −5/-4 dinucleotides of the sgRNA sequence (Supplemental Fig. S7).

### Phenotypic characterization of the mutant library

A major application of mutant libraries is to discover new phenotypes and their causal genotypes. To evaluate this potential in the CRISPR/Cas9 mutant library, we harvested all F1 embryos from 300 randomly selected lines for phenotyping by rearing three broods per line. Apparent morphological differences from the wild-type in terms of body sizes, epidermal coloring, developmental duration, speckle patterns, appendage development, and metamorphosis arrestments were observed among the 33 F1 lines (Fig. 3a, 3b). We correlated these phenotypes with corresponding sgRNA and target site sequencing results, finding individuals from all 33 F1 lines had correct sgRNAs and high editing efficiency at target sites (Supplemental Fig. S8). Many mutant phenotypes were consistent with or theoretically reasonable to previously described mutants, either *B. mori* or other related species, such as *Drosophila*. For example, genes (KWMTBOMO01164, KWMTBOMO03319) of two mutants showed an oily skin phenotype and were previously shown to be involved in transport and accumulation of uric acid in the epidermis (Fujii et al. 2019; Segala et al. 2019), while the developmental arrestment mutant gene, *neverland*, (KWMTBOMO06146) was previously demonstrated to be a key enzyme in ecdysone biosynthesis and caused the arrest of both molting and growth during *Drosophila* development (Yoshiyama et al. 2006; Lang et al. 2012), The wing-deficient mutant gene, *exostosin-1*, was shown to be required for the diffusion of Hedgehog, a key regulator of limb development (Bellaiche et al. 1998). These results indicated that the current library can be used to identify mutants.

**Fig. 3.**
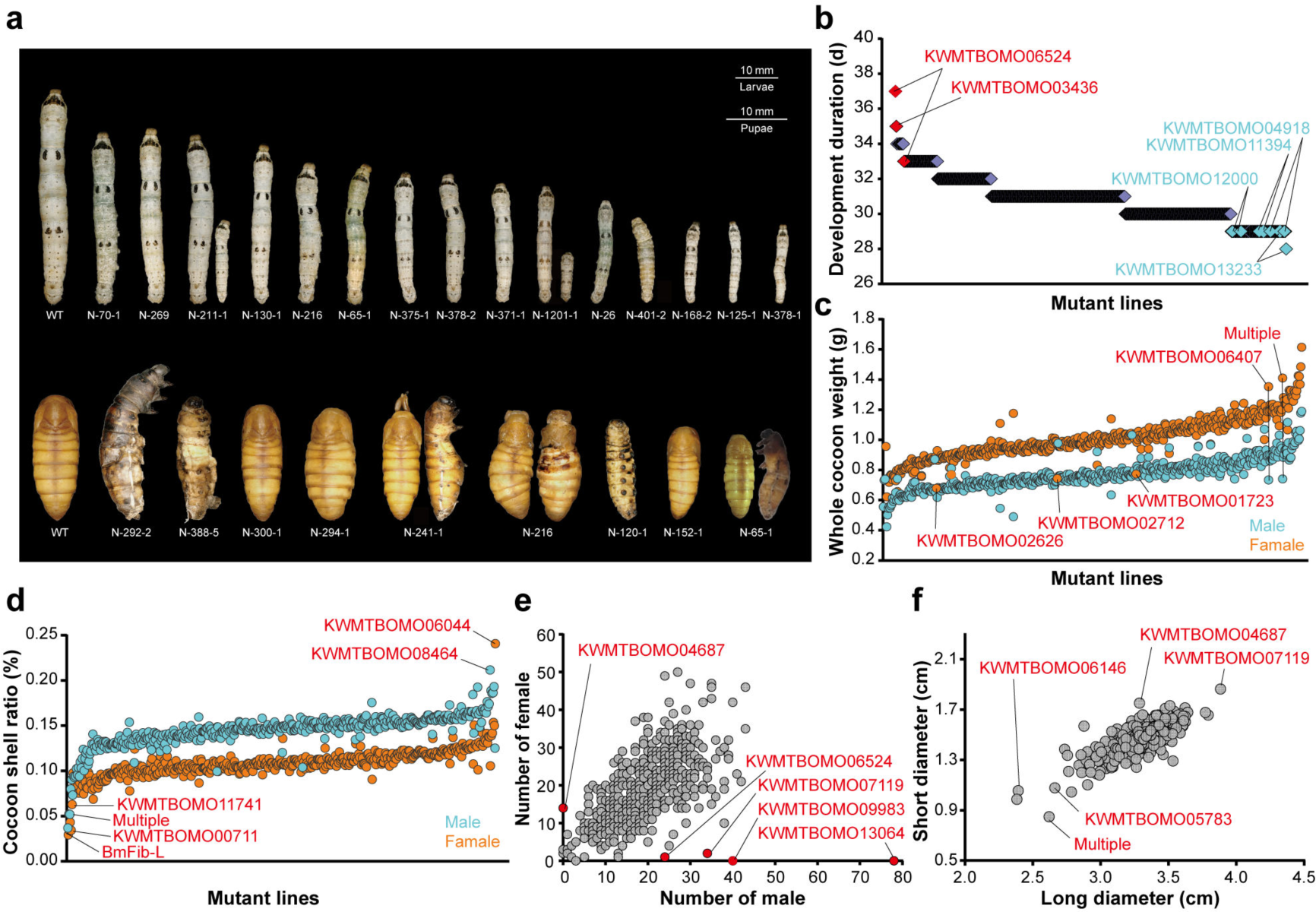
Phenotypic characterization of the mutant library. (a) The body sizes, epidermal coloring, developmental duration, speckle patterns, appendage development, and metamorphosis arrestments of 33 F1 lines. (b) The development duration of different mutant lines. The genes which may be associated with development extending are highlighted by red, those in light blue are more possibly associated with a shortened development. (c) The whole cocoon weight of different mutant lines. The genes possibly associated with whole cocoon weight improvement are highlighted by red. (d) The cocoon shell ratio of different mutant lines. The genes possibly associated with cocoon shell ratio improvement are highlighted by red. (e) The number of males and females in each line. The genes possibly associated with sex differentiation of silkworm are highlighted by red. (f) The short and long diameter of cocoon in each line. The genes that may regulate spinning behaviors are highlighted by red.

As *B. mori* is an economically valuable insect in addition to being a research system, we also investigated the economic traits of the mutant lines. Cocoon weight and shell ratio are the two most obvious traits used to evaluate quality of a commercial strain; we measured the cocoon weight and shell ratio of all 300 mutant lines. Most of these cocoons were average, however we found a few lines that exhibited a notable higher or lower cocoon weight (Fig. 3c) and cocoon shell ratio (Fig. 3d), including a major component of silk protein (*BmFib-L*, Fig. 3d). The multibillion-dollar sericulture industry has long sought an easy way to breed only males, because they produce more silk of a higher quality (Kiuchi et al. 2014; Zhang et al. 2018). We counted the males and females in each line and found that progenies of four lines contained only males while one line only contained females (Fig. 3e), indicating that knocking out these genes might affect the survival of males or females. We also investigated cocoon shapes of these lines (Supplemental Fig. S9), an important trait that affects post-harvesting industrial processes and found several genes that may regulate spinning behaviors (Fig. 3f). These investigations provide targets for strains with improved traits and further illustrate the viability of our library.

### Use of the CRISPR mutant library for pooled screens

Phenotypes of most mutant lines are not visible to the naked eye, but may be responsible for certain physiological changes and exogenous stresses. An important application of the mutant library is pooled screening, which can identify genes responsible for certain fine-designed biotic or abiotic stresses. To explore this potential, we performed a proof-of-principle abiotic stress screening against a common environmental pollutant, cadmium (Cd), exposure to which causes significant economic losses in sericulture areas each year^33^. We selected approximately 100 embryos (1/2 of each brood) from each of the 300 mutant lines and pooled them. After hatching, the larvae were reared on a normal artificial diet in a pooled manner till the 3^rd^ instar. They were then fed an artificial diet containing 30 mg/kg CdCl_2_, which is lethal to the wild type *B. mori*. Surviving silkworms were moved to a fresh artificial diet containing CdCl_2_ to ensure continuous exposure (Fig. 4a). Only 53 individuals survived to the 5^th^ instar. sgRNA and their sequencing revealed that 11 larvae harbored the same sgRNA, targeting KWMTBOMO12902 (Fig. 4b). Sequencing the target region of this sgRNA confirmed highly efficient mutagenesis of KWMTBOMO12902 (Supplemental Fig. S10). Subcellular localization showed that the gene was located in the endoplasmic reticulum and was involved in calcium ion transport. This implies that KWMTBOMO12902 is a Cd stress response gene, and its knockout might cause Cd resistance in silkworms. We checked the performance of this gene in our previous cell line-based genome editing screening results and found that KWMTBOMO12902 was not among the top-ranked positive genes, possibly due to the diversity of Cd ion exposure and responses between individuals and cells, which emphasized the importance of individual mutant pools for finding new strains. We further challenged silkworm individuals and cultured cells with CdCl_2_; we observed that KWMTBOMO12902 was significantly upregulated in the midgut, fat body, and BmE cells of silkworm larvae after Cd exposure (Fig. 4c). An independent CdCl_2_ tolerance test using 3^rd^ instar larvae on a larger scale showed that the mutant line KWMTBOMO12902 had a significantly higher tolerance to CdCl_2_ than the wild type (Fig. 4d). These results indicate that the mutant library in this study could be a powerful resource for identifying new genes responsible for specific physiological functions.

**Fig. 4.**
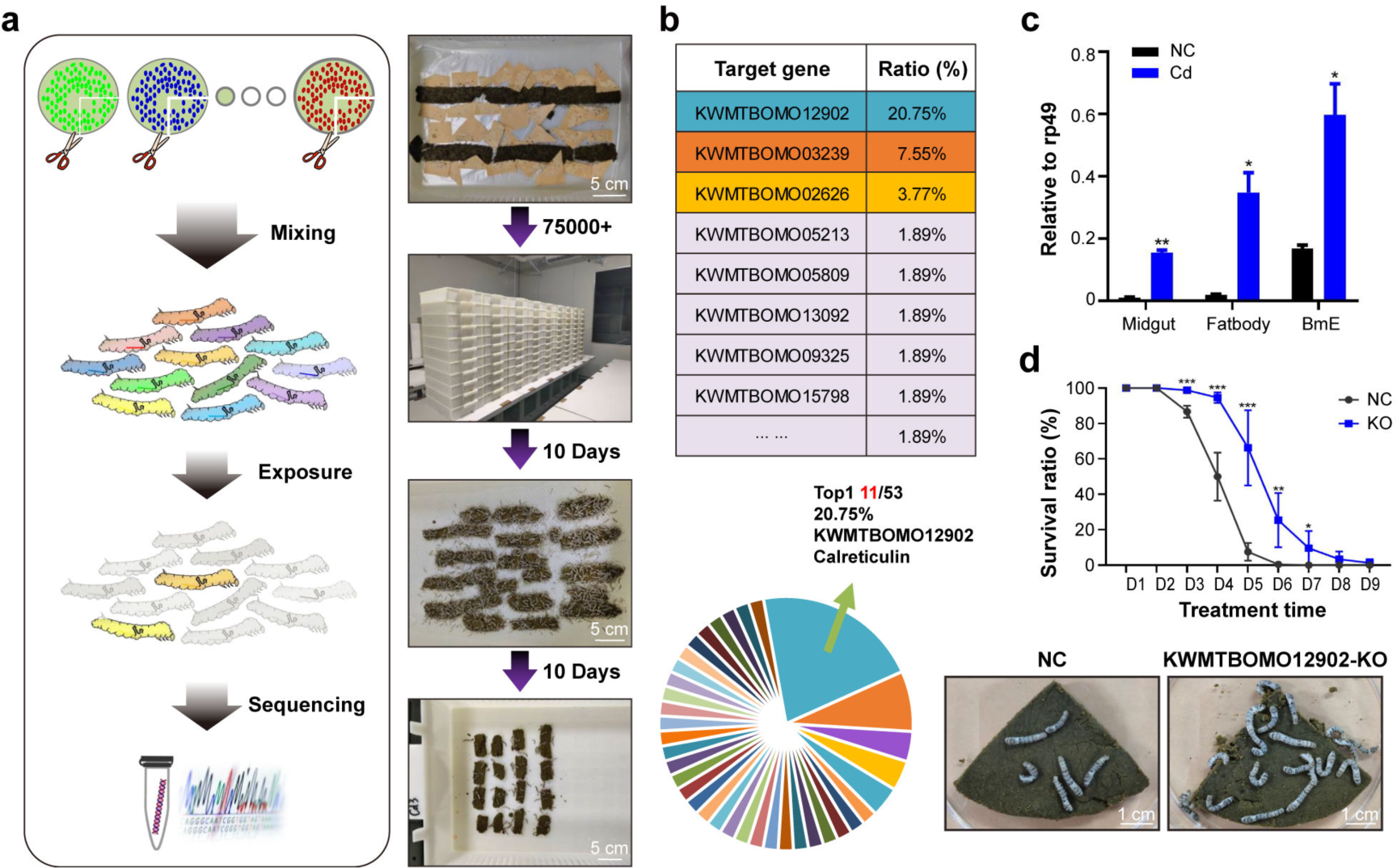
Cadmium exposure screening in the mutant library. (a) The schematic diagram of abiotic stress screening in the mutant library with cadmium exposure (left) and several scenes during the process (right). (b) The target genes survived from persistent cadmium exposure. Individuals with single-guide RNA (sgRNA) targeting KWMTBOMO12902 accounted for the highest proportion. (c) The expression level of KWMTBOMO12902 in midgut, fat body, and BmE cells with or without cadmium exposure. (d) The survival rate of normal strain and KWMTBOMO12902-KO strain in cadmium stress.

## Discussion

Since the early 20^th^ century, the mutant library has been the main source of major breakthroughs in life science innovation and discovery. Several methods, including physical radiation (X-rays, fast neutrons, and gamma-rays), chemical induction (ethyl methane sulphonate and 1,2:3,4-diepoxybutane), random DNA insertion (P-element, *piggy*Bac, and T-DNA), and site-specific mutations (homologous recombination, integrase, RNAi, ZFN, TALEN, and CRISPR/Cas9) have been developed to generate large-scale mutant libraries in a few model organisms such as *Drosophila melanogaster*, *Mus musculus*, *Danio rerio*, *Arabidopsis thaliana*, *Nicotiana tabacum*, *Oryza sativa* L and *Brassica napus*. (Amsterdam et al. 1999; Hrabé de Angelis et al. 2000; Nolan et al. 2000; Gondo, 2008; Lein et al. 2008; Xu et al. 2008; Chang et al. 2012; Haelterman et al. 2014; Lu et al. 2017; Meng et al. 2017; Port et al. 2020; He et al. 2023). Here we used the CRISPR/Cas9 system to construct a large-scale mutant library in a non-model organism, *B. mori*, and demonstrated its application in generating novel mutants with important phenotypic changes, and in identifying new genes with important physiological functions and economic potential, from pooled screening. Research on non-model organisms is crucial for solving complex or specialized biological questions; however, they are severely hindered by the lack of efficient genetic tools and mutant resources. We demonstrated the feasibility of generating a large-scale mutant library using a non-model organism. Furthermore, since we used *piggy*Bac transposon, which can drive gene delivery in a wide range of eukaryotes and has the largest reported cargo capacity (Wu et al. 2007; Chang et al. 2019; Li et al. 2020) as the delivery strategy, the results in this study also provide a straightforward reference for genome-scale mutagenesis in other non-model organisms.

The strategy used in this study has several advantages over previous methods. Traditional mutant *B. mori* variates were collected from either naturally occurring local or chemically/physically-induced mutations (Goldsmith et al. 2005). These have extremely poor phenotype-genotype correlations as the mutations occur randomly across the genomic region, and different mutations have diverse genetic backgrounds (Tong et al. 2022). It usually takes years to map a certain phenotype to its causal genetic alterations (Liu et al. 2010; Atsumi et al. 2012; Yamaguchi et al. 2013). In *Drosophila* spp. and mice, large-scale mutagenesis can also be achieved through subsequently developed targeted methods such as homologous recombination and RNAi, but both are extremely inefficient in *B. mori* and many other non-model organisms. The CRISPR technology used in our study overcame all the shortcomings of previous methods. First, all mutagenesis was guided by the sgRNA sequences, which were designed to target only the coding regions of the genome, and all mutant lines in the library were truly functionally mutated. Second, the mutated gene and mutant sequences in each line can be tracked through integrated sgRNA sequences by simple targeted PCR amplification and sequencing, meaning all mutant lines in the library have clearer and closer genotype-phenotype correlations than previous resources. Third, as demonstrated, the efficiency of the CRISPR library for generating large-scale mutants is sufficient for any laboratory with common equipment. Our study also provided many parameters that may further improve this efficiency, such as the high input/low input strategy to avoid repeated sgRNA lines, the backcrossing strategy to reduce multiple sgRNA insertions, and the binary vector system to avoid loss of growth-deficient lines with lethal genes.

*B. mori* is one of the most economically beneficial insects and a representative lepidopteran insect because of its crucial role in producing the best-known animal fibers for thousands of years, and its finely decoded genomic sequences (Xia et al. 2014; Tong et al. 2022). Systematic functional investigation of *B. mori* genes is required for both fundamental research and agricultural applications. To achieve this goal, we established multiple genetic manipulation tools, including transgenesis, site-specific recombinases, ZFN, TALEN, and CRISPR/Cas9, as well as a high-throughput CRISPR library screening platform in cultured *B. mori* cells (Ma et al. 2012; Ma et al. 2013; Duan et al. 2013; Liu et al. 2014; Ma et al. 2014b; Li et al. 2018; Chang et al. 2020). However, because the accumulation of genetic resources and research in *B. mori* relies heavily on the study of orthologous genes in related model organisms, the function of more than 80% of the *B. mori* genes remains unclear. We herein present the first high-throughput platform to generate large-scale whole-animal-level mutant libraries and provide a hypothesis-free, cost-effective pipeline to rapidly investigate the function of *B. mori* genes on a genome-wide scale. As a proof-of-principle demonstration, we designed and constructed a plasmid library containing 92,917 sgRNAs targeting 14,645 protein coding genes, with high-quality coverage, distribution, and accuracy. A total of 1726 sgRNA lines were generated by microinjection of over 60,000 embryos, and 300 mutant lines with deciphered mutant sequences containing over 30 obvious phenotypic changes were obtained. These resources provide a powerful platform for functional annotation of *B. mori* genomes.

We previously established a pooled loss-of-function screening platform in cultured *B. mori* cells and demonstrated its effectiveness in identifying genes involved in multiple biotic and abiotic stresses (Chang et al. 2020; Liu Y et al. 2021). It remains a significant challenge to perform pooled genetic screening at the whole-animal level due to the lack of mutant resources and an efficient screening method. Here we performed the first in vivo pooled genetic screening of *B. mori*. We showed that the CRISPR mutant library was highly efficient in identifying the genes responsible for stress. The screening results were mutually complemented genome-wide CRISPR screenings in cultured cells, and the selected mutants showed high performance when checked individually. We further identified the KWMTBOMO12902 gene, the absence of which significantly increased the tolerance to Cd exposure. Cd exists extensively in the ecosystem as a typical pollutant. It has been identified as a global threat because it cannot be degraded or resolved, is gradually enriched up the biological chain, and is severely harmful in various animals and plants, including humans (Yang et al. 2022). Studies aiming to decipher signaling and metabolic pathways involved in the pathological effects associated with Cd exposure are urgently needed, but it is difficult to perform large-scale toxicant exposure experiments in mammals. Here we used *B. mori* for large-scale pooled screens, and with the nature of *B. mori* being sensitive to environmental pollutants, it provided a potential target to explore organismal detoxification mechanisms (Mao et al. 2019). Environmental Cd exposure causes significant economic damage to the sericulture industry. Results showed the KWMTBOMO12902 gene mutant exhibited a high tolerance to Cd, even at a dosage much higher than that of environmental Cd, suggesting that KWMTBOMO12902 can be used to breed a Cd-tolerant silkworm strain.

Although the CRISPR/Cas9 system has been a powerful genome editing tool for more than 10 years, designing sgRNAs with high activity remains a major challenge (Chen and Wang 2022). As has been recognized in a number of previous reports and in the current study, not all sgRNAs can cleave their target DNA, and the cleavage efficiencies of different sgRNAs vary significantly (Li et al. 2022). A key parameter that determines the efficiency of sgRNA is its sequence and structural properties. We systematically investigated the effects of sgRNA sequence composition by sequencing 1436 mutant silkworms and found that the GC content in the 2–10 nt of the sgRNA (distal to the PAM) region, C content within the PAM sequences, G at the 3′ end, G at the 5′ end, and G/CG/C dinucleotides within the seed region are beneficial for efficient editing. We also investigated the effect of the local chromosome state on editing efficiency and found that 6mA methylation at the target site has a positive effect. These results provide practical guidance for the design of sgRNA for a successful genome editing experiment in *B. mori* and increase the understanding of the heterogeneity of sgRNA functions.

In conclusion, we have established the first genome-scale CRISPR mutagenesis platform at the whole-animal level in a non-model multicellular organism. A plasmid library containing 92,917 sgRNA-expressing vectors and a collection of 1726 sgRNA transgenic lines was established. By generating and analyzing 300 CRISPR mutant lines, we showed that the CRISPR library is a powerful resource for the rapid discovery of unknown gene functions and for investigating long-standing questions in silkworms and entomology. Applying the established mutant lines, we discovered and highlighted a large set of new genes that can cause divergent visible phenotypic changes, and several genes that have promising potential applications in breeding strains, with higher yield or greater tolerance for the sericulture industry. These resources are preserved in the Biological Research Center of Southwest University and are shared worldwide through the silk genome database, SilkDB 3.0 (Lu et al. 2020). Furthermore, the number of sgRNA transgenic and mutant lines will continue to increase. We believe that this study lays the foundation for large-scale functional genomics of *B. mori* and the rapid identification of novel genes involved in diverse biological processes. More importantly, the present study also sheds light on the application of genome-scale mutagenesis to other non-model organisms.

## Methods

### Silkworm strains and rearing conditions

Two silkworm strains, Dazao and Nistari, were used in the present study. Dazao was used to inject the sgRNA library, and Nistari to express Cas9. Dazao was reared in our laboratory while Nistari was provided by Prof. Anjiang Tan from Jiangsu University of Science and Technology, Zhenjiang, China. The silkworms were fed fresh mulberry leaves or artificial diet at 25 ℃ and 80 % relative humidity.

### Design and construction of the vectors

The *piggy*Bac transposase expression vector A3-helper was obtained from stocks in our laboratory. pUC57-3Xp3-EGFP-sv40 was constructed by replacing DsRed in pUC57-3Xp3-DsRed-sv40 (3DS) (Ma et al. 2014) with the EGFP sequence. pUC57-U6-gRNA (*Aar*I) was constructed by replacing the *Bbs*I restriction site in pUC57-gRNA (Ma et al. 2014) with an *Aar*I restriction site. PUC57-3Xp3-EGFP-sv40 was cloned into the *Age*I/*Asc*I sites of the pBac-modified vector to generate pB-modified {3Xp3-EGFP-sv40}. pUC57-U6-gRNA(*Aar*I) was cloned into the *Nhe*I/*Asc*I site of pB-modified {3Xp3-EGFP-sv40} to generate the vector pB-modified {3Xp3-EGFP-sv40}{U6-gRNA(*Aar*I)}, named p200.

### sgRNA library design and synthesis

sgRNAs were designed using the CCTop method (Aach et al. 2014). All sgRNAs selected had 5′ G added to improve the U6 transcription efficiency. The sgRNA library was encoded within 70-nt oligonucleotides and synthesized on 92,917 arrays using the services of the Beijing Genomics Institute (China). The oligonucleotide sequence used was 5’-TACAA AATAT CGTGC TCTAC AAGTG NNNNN NNNNN NNNNN NNNNN GTTTT AGAGC TAGAA ATAGC AAGTT-3’.

### Construction of sgRNA library

The DGP-1 fragment (U6 promoter) was PCR-amplified from p200 using the primers KU-1R and DG-1R. The pool of sgRNA oligonucleotides was amplified by PCR using primers DP-2F and DP-2R and named DGP-2 fragments (multiple sgRNAs). The DGP-3 fragment (sgRNA scaffolds) was amplified by PCR from p200 using the primers DG-3F and KU-1F. These three fragments were linked using overlap PCR to generate DGP-4 fragments (U6-sgRNA library). All PCR were performed using PrimeSTAR Max DNA polymerase (Takara, Japan). The DGP-4 fragments were then cloned into the AscI/NheI (NEB, USA) site of the p200 vector to construct the sgRNA library (p200 library) according to the following protocol: The gel-purified DGP-4 fragments (200 ng) and p200 vector (200 ng) were mixed and ligated in a 50 µL T4 ligation reaction (NEB, USA) according to the manufacturer’s instructions. The ligation reaction (1 µL) was transformed into 50 µL of *Escherichia coli* HST08 premium electro-cells (Takara, Japan) using a Gene Pulser Xcell (Bio-Rad, USA) according to the manufacturer’s instructions. To ensure the diversity of the sgRNA library, 15 parallel electro-transformations were performed with ampicillin selection, which resulted in >1,000× library coverage. All clones were collected, and the plasmid library was extracted using a QIAprep Spin Miniprep Kit (Qiagen, Germany).

### Microinjection and positive transgenic screening

After mating, the female moths were placed on paper starched with paste to lay eggs in the dark. Eggs were collected every 0.5 h and arranged on slides (approximately 100 eggs per slide). After the eggs were fixed on slides, plasmids were injected using a microinjection apparatus (Eppendorf AG, Germany). The injection was completed within 60–220 min after laying. The injected plasmid was mixed with sgRNA plasmid: Helper plasmid = 1 × 1 (molar ratio of the substance), and the final concentration was 400 ng/μL. Plasmids (10–15 nL) were injected into each egg. Immediately after injection, the eggs were sealed with non-toxic glue (Instant Strong Glue Mini; Japan). The eggs were maintained at a relative humidity above 85% until they hatched. After hatching, the larvae were reared according to the type of vector injected. The G0 generation obtained by self-cross or backcross seed production were detected and screened using a macrosteric fluorescence microscope (OLYMPUS DP73, Japan) and fed separately.

### Pooled genetic screening

After collecting the eggs from the silkworm mutant library, approximately 1/2 of each brood was gathered from every mutant line, totaling approximately 30,000–50,000 eggs. Subsequently, these larvae were uniformly hatched and fed a normal artificial diet until the 3^rd^ instar. The larvae were fed an artificial diet containing 30 mg/kg CdCl_2_. Subsequently, dead larvae were removed, and survival was supported with a fresh diet. Ultimately only 53 individuals survived until the 5^th^ instar. They were used as donors for the extraction of genomic DNA and sgRNA sequences.

### sgRNA sequencing, target sites sequencing, and data analysis

The genome was amplified using primers with distinct barcodes and sequenced using Illumina sequencing technology. Low-quality raw sequencing data of amplicons were filtered out using Trimmomatic (v.0.39) (Bolger et al. 2014) (LEADING:3 TRAILING:3 SLIDINGWINDOW:4:15 MINLEN:50). Paired-ended clean reads were merged using FLASH (v.1.2.11) (Magoč et al. 2011). Cutadapt (v.1.18) (Marcel et al. 2011) was used to split the data based on the unique barcode of each sample. Bowtie2 (v.2.2.5) (Langmead et al. 2012) was used for aligning to identify the sgRNA sequences in each sequenced sample. Based on the detected sgRNA, Primer3 (v.2.4.0) (Untergasser et al. 2012) was used to design primers for each target site, amplified the genomes of different samples using diverse primers with specific barcodes, and conducted second-generation sequencing. Barcode sequences associated with each sample are listed in Supplemental Table S2. The analysis methods for filtering out, merging, and splitting of target site sequences were the same as those used in the sgRNA data analysis, while mutation detection of target sites was performed using CRISPResso (v.2.0.45) (Clement et al. 2019).

### Cd exposure experiment

To further determine the function of KWMTBOMO12902 in Cd tolerance, we separately treated the KWMTBOMO12902-KO silkworm strain with Cd exposure. Embryos were hatched and fed a fresh artificial diet. In the 3^rd^ instar, the artificial diet was supplemented with 30 mg/kg CdCl_2_. The survival rate was calculated daily. To further investigate the role of KWMTBOMO12902 in Cd tolerance, we exposed KWMTBOMO12902-KO strain silkworms to Cd. Embryos were hatched and fed a fresh artificial diet. In the 3^rd^ instar, the artificial diet was supplemented with 30 mg/kg CdCl_2_. The survival rate was calculated daily.

### Phenotypic characterization

Fourth-instar larvae of 300 strains of the F1 generation were screened using a stereo-fluorescence microscope (OLYMPUS DP73, Japan), and all double-positive individuals (the eye with green light and the body with green light) in each brood were selected for subsequent research. Mutants with significant developmental abnormalities (larval development speed and individual size, larval pattern, pupation process, cocoon color and type, and moth-changing process) were selected. The cocoon weight and pupal weight of all double-positive males and females in each brood were measured separately using an analytical balance, and the sex ratio was counted simultaneously. The cocoon shape was measured for length and breadth using ImageJ software (https://imagej.nih.gov/ij/index.html), and 10 female and 10 male cocoons were randomly measured.

### Quantitative real-time polymerase chain reaction (qPCR)

The larvae were dissected to obtain different tissues, and the total RNA from BmE cells or different tissues was extracted. For each sample, 1 μg of RNA was subjected to reverse transcription using the one-step gDNA removal and cDNA synthesis SuperMix Kit (TransGen Biotech, China). qPCR was performed using SYBR® Premix Ex Taq™ II (Takara, Japan), and target gene expression was normalized to the reference gene rp49. The test was performed independently at least three times. The primer sequences for the target genes were as follows: KWMTBOMO12902-qPCR-F: TGAAGAGCGAACAAGACGAG; KWMTBOMO12902-qPCR-R: TTCCTCATCGTCAGGCTTTT; Rp49-qPCR-F CAGGCGGTTCAAGGGTCAATAC; and Rp49-qPCR-R TGCTGGGCTCTTTCCACGA.

## Data access

The raw data generated in this study were submitted to the NCBI BioProject database (https://www.ncbi.nlm.nih.gov/sra) under accession number PRJNA901072.

## Acknowledgments and funding sources

The authors would express heartfelt thanks to Dr. Feng Wang, Dr. Zhangchuan Peng, Dr. Wenliang Qian, Dr. Kaiyu Guo, Dr. Qingsong Liu, Miss Xia Zhang, and Miss Xiaoxu Chen for their kind help with the microinjection experiments. This work was supported by grants from the National Natural Science Foundation of China (No. 32122084), the Chongqing Natural Science Foundation (No. cstc2021ycjh-bgzxm0005), PhD Start-up Foundation of Southwest University (no. SWU120012), and Fundamental Research Funds for Central Universities (No. SWU-KT22042). None of these funding sources played a role in the design of the study, collection, analysis, and interpretation of data, or in writing the manuscript.

## Author Contributions

SM conceived of the study and designed the experiments. TZ designed the sgRNA and performed all bioinformatic analyses. RW constructed the plasmid library. YL performed the pooled screening experiments. PW, LJ, and AW generated and maintained the transgenic lines with the help of JC, LS, HS, RS, WL, DL, NZ, WH, XW, and WX. XL performed the genotyping experiments. LJ and SM wrote the manuscript with the help of QX. All authors have read and approved the final manuscript.

